# In Silico Screening of Indian Medicinal Herb Compounds for Intestinal α-Glucosidase Inhibition with ADMET and Toxicity Assessment for Postprandial Glucose Management in Type-2 Diabetes

**DOI:** 10.64898/2026.03.01.708840

**Authors:** Ashim Chandra Roy, Ilora Ghosh

**Affiliations:** School of Environmental Sciences, Jawaharlal Nehru University, New Delhi-110067, India

**Keywords:** Hyperglycemia, Ashwagandha, Tea, Curcumin, Glucose absorption

## Abstract

Postprandial hyperglycemia is a major concern in type 2 diabetes, and inhibition of intestinal α-glucosidases [maltase–glucoamylase (MGAM)] is an established method for controlling post-meal glucose excursions. In this study, we conducted an in-silico screening of phytochemicals from different well-known medicinal plants (*Withania somnifera, Rauwolfia serpentina, Curcuma longa, and Camellia* sinensis) against MGAM, using the clinically approved inhibitor miglitol as reference for docking protocol validation. Molecular docking revealed that miglitol binds to MGAM with a binding energy of −6.86kcal/mol and an RMSD of 1.04 (with co-crystal structure; PBD ID:3L4W); however, several phytochemicals exhibited binding affinities equal to or stronger than miglitol. Among these, Withanolide B (−9.25kcal/mol) and Withanone (−7.57kcal/mol) from *Withania somnifera* showed the highest predicted affinities, indicating robust engagement of the MGAM catalytic pocket. *Rauwolfia serpentina* alkaloids such as yohimbine (−8.50kcal/mol) and raubasine (−8.46kcal/mol) also displayed strong binding energies, whereas curcuminoids (curcumin −6.36kcal/mol; deoxycurcumin −6.35kcal/mol) and tea catechins (e.g., epicatechin gallate −6.85 kcal/mol) demonstrated moderate affinity. Interaction analysis showed that top-ranking compounds formed extensive hydrogen-bonding and hydrophobic interactions with key catalytic residues of MGAM, suggesting stable occupancy of the active site. In-silico ADME profiling predicted favorable gastrointestinal absorption for lead phytochemicals, supporting their potential for oral intestinal action. Collectively, these results identify plant-derived ligands with binding energies comparable to or exceeding that of miglitol, highlighting *Withania somnifera* withanolides as priority candidates for experimental validation in enzyme inhibition assays and glucose tolerance models, and providing a focused set of natural MGAM inhibitors for further translational investigation in postprandial glucose control.

## Introduction

Controlling postprandial glucose (PPG) is an essential clinical target in type-2 diabetes mellitus (T2DM) because spikes in glucose after meals contribute significantly to overall glycemic burden and are associated with diabetes-related complications such as cardiovascular disease and endothelial dysfunction, making this control crucial for improved outcomes (α-glucosidase inhibitors improve metabolic profiles and can reduce long-term complications of hyperglycemia)(Akmal et al. 2025). Elevated postprandial glucose plays a central role in the pathogenesis of diabetes-associated vascular damage and chronic inflammation, emphasizing the importance of targeting glucose excursions after meals (Hossain et al. 2020).Reducing PPG reduces glucose excursions and improves markers of glycemic control, indicating its value beyond fasting glucose levels alone. α-Glucosidase inhibitors (AGIs) slow carbohydrate digestion in the small intestine by delaying the breakdown of complex carbohydrates into absorbable glucose, thus attenuating postprandial glucose peaks (Han et al. 2025).AGIs such as acarbose and miglitol are established therapeutic options to blunt PPG in patients with T2DM, and evidence from clinical meta-analyses and systematic reviews supports their use due to reductions in postprandial glucose and insulin responses after carbohydrate loads (Alssema et al. 2021; Van de Laar et al. 2005). The mechanism of α-glucosidase inhibition, that prevents the conversion of oligo- and disaccharides to monosaccharides and ultimately delays glucose absorption from the gut, distributing glucose delivery into the bloodstream more gradually and reducing early postprandial peaks (Han et al. 2025). Although synthetic AGIs are effective, gastrointestinal side effects such as flatulence and abdominal discomfort can limit tolerability and adherence in some patients(Akmal et al. 2025). Because of this, research has focused on natural α-glucosidase inhibitors from plant sources, which may offer similar benefits with fewer adverse effects and additional bioactivities (Dirir et al. 2022). Phytochemicals such as flavonoids, terpenoids, phenolic acids, anthocyanins, tannins, and procyanidins have been widely reviewed for their potential α-glucosidase inhibitory activity and their capacity to control postprandial hyperglycemia in vitro (Hossain et al. 2020).Compounds like Taxumariene F, Morusin, and Procyanidin A2 isolated from diverse plant species have shown potent inhibitory activity against α-glucosidase comparable to synthetic standards in biochemical studies (Dirir et al. 2022).Traditional herbs including *Gymnema sylvestre, Momordica charantia, Trigonella foenum-graecum, Tinospora cordifolia,* and *Curcuma longa* have been documented for antidiabetic potential and content of bioactive phytoconstituents that may contribute to glycemic regulation(Gaonkar and Hullatti 2020). Phytochemical screening of various plant extracts has identified botanical sources with significant α-glucosidase inhibitory activity, suggesting that these natural products may be promising leads for further computational docking and drug design (Mujawdiya and Kapur 2020). Virtual screening and molecular docking studies of plant-derived compounds have become essential tools in accelerating the identification of high-affinity ligands targeting intestinal α-glucosidase before experimental validation. ADMET profiling and toxicity assessment complement docking by evaluating drug-like properties and safety, which are critical in selecting compounds with translational potential (Dirir et al. 2022). In summary, controlling postprandial glucose through intestinal α-glucosidase inhibition is a validated therapeutic strategy in T2DM, and phytochemicals from traditional medicinal herbs represent a rich source of natural inhibitors that warrant further computational and experimental investigation (Hossain et al. 2020).

Although many in vitro and in vivo studies report glucose-lowering effects of crude plant extracts, these studies often lack precision because they do not identify the specific molecules responsible for the activity. As a result, it is difficult to establish clear structure–activity relationships or advance from extract-based observations to well-defined, optimizable drug leads that act on specific targets such as intestinal α-glucosidase. In silico molecular docking and computational ADMET & toxicity screening address this limitation by predicting how individual phytochemicals bind to a target enzyme, estimating binding affinity, and evaluating pharmacokinetic and safety profiles early in the discovery pipeline, thereby enabling efficient prioritization of specific compounds for further experimental validation and reducing the cost and time of drug development processes that would otherwise rely on trial-and-error testing of crude extracts(Agu et al. 2023; Asiamah et al. 2023).

In this study, we selected phytochemicals from *Rauwolfia serpentina*, *Curcuma longa*, *Camellia sinensis,* and *Withania somnifera* based on their ethno-pharmacological use and reported bioactivity related to glucose metabolism. *Rauwolfia serpentina* extracts contain diverse alkaloids and phenolic compounds with documented hypoglycemic activity (Azmi and Qureshi 2012; Kavitha et al. 2025). Extracts of *Curcuma longa* (turmeric) contain curcuminoids and phenolics with reported antidiabetic potential (Lekshmi et al. 2014; Okechukwu et al. 2023). *Camellia sinensis* is rich in catechins and flavonoids associated with antidiabetic effects (Ansari et al. 2022; Brimson et al. 2022). *Withania somnifera* (*Withania somnifera*) contains withanolides and related bioactive compounds with suggested roles in glucose metabolism modulation (Alhasani et al. 2026).

Despite the wealth of extract-based research, it remains unclear which specific phytochemicals play a significant role in hypoglycemic activity, indicating a critical gap in the systematic identification, ranking, and screening of individual bioactive compounds. By applying basic in silico techniques, including molecular docking to predict binding interactions with the target protein, followed by ADMET and toxicity profiling to assess pharmacokinetic behavior and safety, this research gap can be addressed, providing a valuable boost to the pharmaceutical industry for target-based drug discovery. The present study aims to prioritize phytochemical leads with strong inhibitory potential against intestinal α-glucosidase, a key protein involved in glucose absorption, along with favorable drug-like properties. This approach provides a focused set of candidates for future experimental validation and accelerates the discovery of novel, safer, plant-derived α-glucosidase inhibitors for postprandial glucose control in type 2 diabetes.

## Material and Methods

### Phytochemical Data Collection

Phytochemical constituents of *Rauwolfia serpentina*, *Curcuma longa, Camellia sinensis, Tupistra nutans,* and *Withania somnifera* were compiled through targeted literature searches in PubMed using scientific plant names combined with terms such as “phytochemical,” “chemical constituents,” and “secondary metabolites.” Identified compound names were standardized and cross-referenced with PubChem to obtain canonical SMILES, InChIKeys, and structural data. Additionally, phytochemical occurrence information was verified and supplemented using the IMPPAT (Indian Medicinal Plants, Phytochemistry and Therapeutics) database at https://cb.imsc.res.in/imppat/ and Dr. Duke’s Phytochemical and Ethnobotanical Databases at https://phytochem.nal.usda.gov/search to ensure comprehensive coverage of reported plant compounds.

### In Silico Screening Workflow

The curated phytochemical library was prepared for docking by converting canonical SMILES into appropriate 3D conformations. The intestinal α-glucosidase receptor, maltase-glucoamylase (MGAM), was selected for molecular docking, and the crystal structure of its N-terminal catalytic domain complexed with miglitol (PDB ID: 3L4W) was retrieved from the RCSB Protein Data Bank (https://www.rcsb.org/structure/3l4w) for structural modeling and docking studies. Protein preparation involved removal of crystallographic water molecules, addition of polar hydrogens, and optimization of geometry prior to docking. Phytochemical ligands were prepared by optimizing torsional flexibility and aromatic conformations. Docking results were ranked based on binding affinity, and top-ranked compounds were selected for pharmacokinetic and toxicity evaluation using predicted ADMET and toxicity profiles from established in silico tools.

### In Silico Molecular Docking Analysis

Phytochemicals were docked against human maltase-glucoamylase (MGAM), an intestinal α-glucosidase involved in glucose release during carbohydrate digestion. The crystal structure of the N-terminal catalytic domain of human maltase-glucoamylase in complex with miglitol (PDB ID: 3L4W) was retrieved from the RCSB Protein Data Bank (RCSB PDB:3L4W; Crystal complex of N-terminal Human Maltase-Glucoamylase with miglitol). Protein preparation involved removing crystallographic water molecules, adding polar hydrogens, and assigning Kollman charges. Ligand structures were optimized for torsional flexibility and aromaticity. Molecular docking was performed using AutoDock4, where the docking grid was centered on the miglitol-binding site of MGAM with coordinates x = 45.38, y = 92.35, z = 35.82, a grid spacing of 0.450 Å, and grid dimensions of 60 × 60 × 60 points. The Genetic Algorithm was applied with a maximum of 2.5 × 10□ energy evaluations to predict optimal ligand–protein binding conformations. Before docking the test phytochemicals, the docking protocol was validated by re-docking the standard inhibitor miglitol into the MGAM active site. Accuracy was assessed by calculating the root-mean-square deviation (RMSD) between the crystallographic and re-docked poses using the DockRMSD platform (https://aideepmed.com/DockRMSD/). An RMSD value ≤ 2.0 Å was considered acceptable, and only after successful protocol validation were the selected phytochemicals docked using the same parameters.

### ADMET Analysis

Phytochemicals were subjected to ADMET (Absorption, Distribution, Metabolism, Excretion, and Toxicity) evaluation using in silico prediction tools to assess their pharmacokinetic and drug-likeness profiles. Canonical SMILES of docked phytochemicals were uploaded to SwissADME to predict key ADMET parameters, including oral bioavailability, human intestinal absorption, blood-brain barrier permeability, cytochrome P450 enzyme interactions, and clearance properties. Predicted data were used to identify compounds with favorable pharmacokinetic attributes suitable for further consideration as potential therapeutic leads.

### Toxicology Analysis

The top phytochemicals were evaluated for toxicity profiles using in silico toxicity prediction platforms ProTox 3.0 (http://tox.charite.de/protox3/index.php) to estimate potential adverse effects. Parameters assessed included acute toxicity (LD□□), hepatotoxicity, cardiotoxicity (hERG inhibition risk), mutagenicity, carcinogenicity, and skin sensitization potential.

## Results

### Docking Protocol Validation

To validate the docking protocol, the co-crystallized ligand was extracted and re-docked into the same binding site using identical grid and scoring parameters. In the crystal structure, the ligand is deeply embedded in the pocket, forming strong hydrogen bonds with ASP327, ASP443, ASP542, ARG526, and HIS600, and engaging in hydrophobic/π contacts with TRP406, TRP441, PHE575, MET444, as well as additional contacts with ILE328, ILE364, and TYR299 **(Figure 1A)**. The redocked pose recapitulates the overall orientation and retains the majority of these interactions, with 10 of 12 interacting residues conserved (∼83 %) with a binding energy of −6.85 kcal/mol and an RMSD of 1.048 Å (DockRMSD) relative to the crystal pose, demonstrating high fidelity of the docking protocol **(Figure 1B)**. Minor deviations are observed in flexible regions, including an unfavorable acceptor–acceptor contact with TRP539 not seen in the crystal, and slight geometric perturbations of some hydrogen bonds. These discrepancies likely arise from receptor rigidity and scoring limitations, but do not compromise the core interaction network, confirming the reliability of the docking setup for subsequent ligand evaluation.

**Figure 1.**
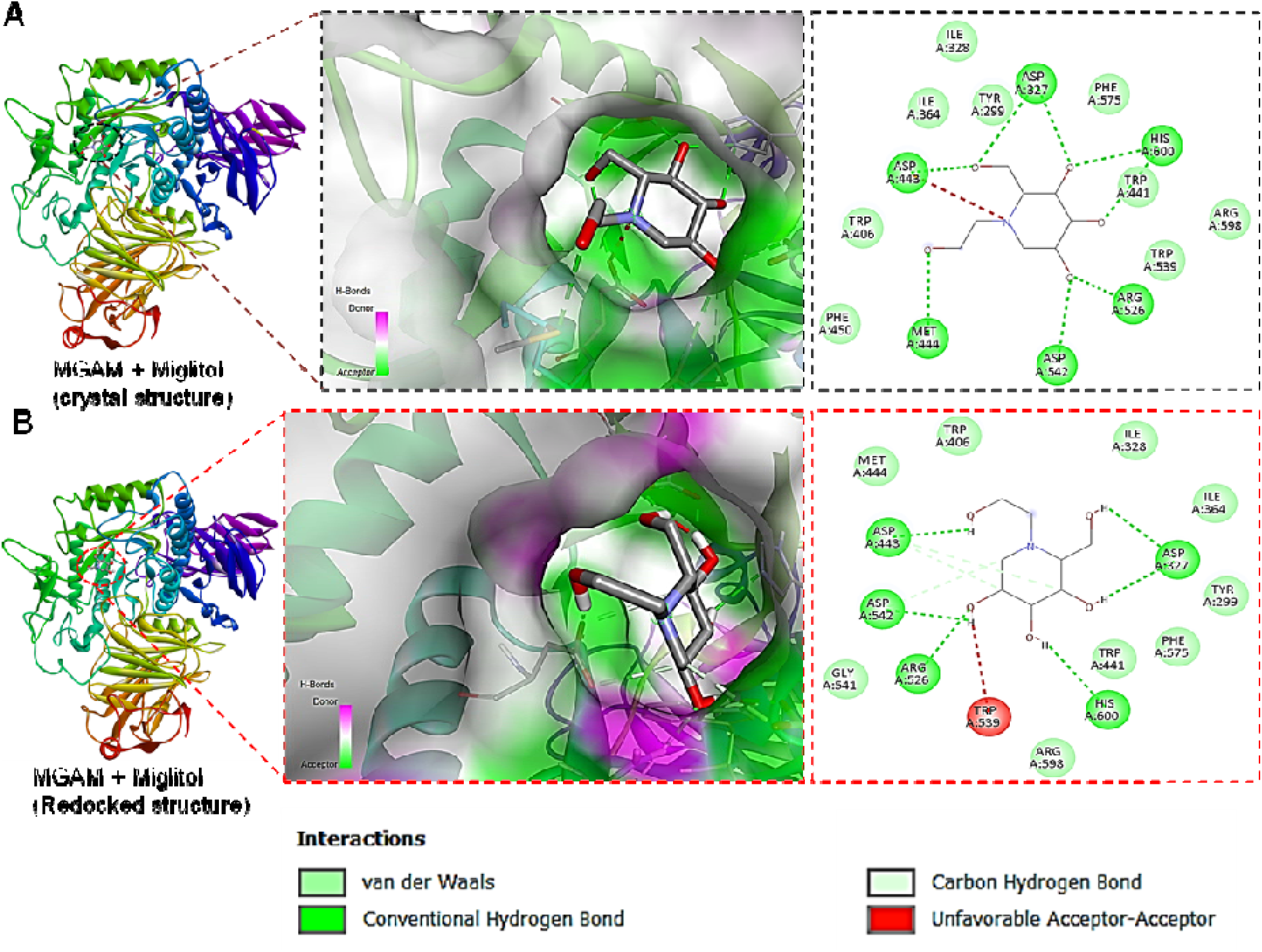

### In silico α-glucosidase inhibition activity of *Rauwolfia serpentina* phytochemicals

Molecular docking analysis was carried out to investigate the α-glucosidase inhibitory potential of five major phytochemicals from *Rauwolfia serpentine* including, raubasine, yohimbine, serpentine, reserpine, and ajmaline against MGAM. All compounds were successfully accommodated within the MGAM catalytic pocket and exhibited favorable binding energies ranging from −6.68 to −8.50 kcal/mol, reflecting variable but meaningful interaction strengths **(Figure 2A–E)**. Among these phytochemicals, raubasine showed a strong binding affinity toward MGAM with a binding energy of −8.46 kcal/mol (**Figure 2A**). The ligand established multiple hydrogen bonds with key catalytic residues, including Asp542, Asp443, Tyr214, Thr204, and Thr205, which are crucial for enzymatic activity. These interactions were further strengthened by π–π and hydrophobic contacts with residues such as Tyr299, Trp406, Phe450, Met444, and Ile328, indicating stable occupancy of the active site **(Figure 2A)**. Yohimbine exhibited the highest binding affinity among the tested phytochemicals, with a binding energy of −8.50 kcal/mol **(Figure 2B)**. It formed hydrogen bonds with Asp542, Asp203, Thr204, and Thr205, while extensive π–π and hydrophobic interactions with Tyr299, Trp406, Phe450, Met444, and Asp443 contributed to enhanced stabilization within the catalytic pocket. Serpentine demonstrated favorable binding with a binding energy of −7.62 kcal/mol **(Figure 2C)**. The compound interacted with MGAM through hydrogen bonds involving Thr205, Thr204, Asp203, and Tyr214, along with supportive hydrophobic and aromatic interactions with Tyr299, Trp406, Met444, Asp443, Phe450, and Arg526, collectively stabilizing the ligand–enzyme complex. Ajmaline also showed strong binding affinity (−7.53 kcal/mol) and a well-defined interaction pattern within the MGAM active site **(Figure 2E)**. Hydrogen bond interactions with Asp203, Asp443, and Asp542 were complemented by hydrophobic and π–π interactions involving Tyr299, Trp406, Met444, Phe450, Ile328, and Ile364, suggesting effective engagement of the catalytic pocket. In contrast, reserpine displayed comparatively weaker binding affinity (−6.68 kcal/mol) **(Figure 2D)**. Although it formed hydrogen bonds with Gln603 and Gly602 and established π–π and hydrophobic interactions with Trp406, Phe450, Tyr299, Phe575, and Gly604, the overall interaction network was less extensive, indicating reduced stability within the active site relative to the other alkaloids.

**Figure 2.**
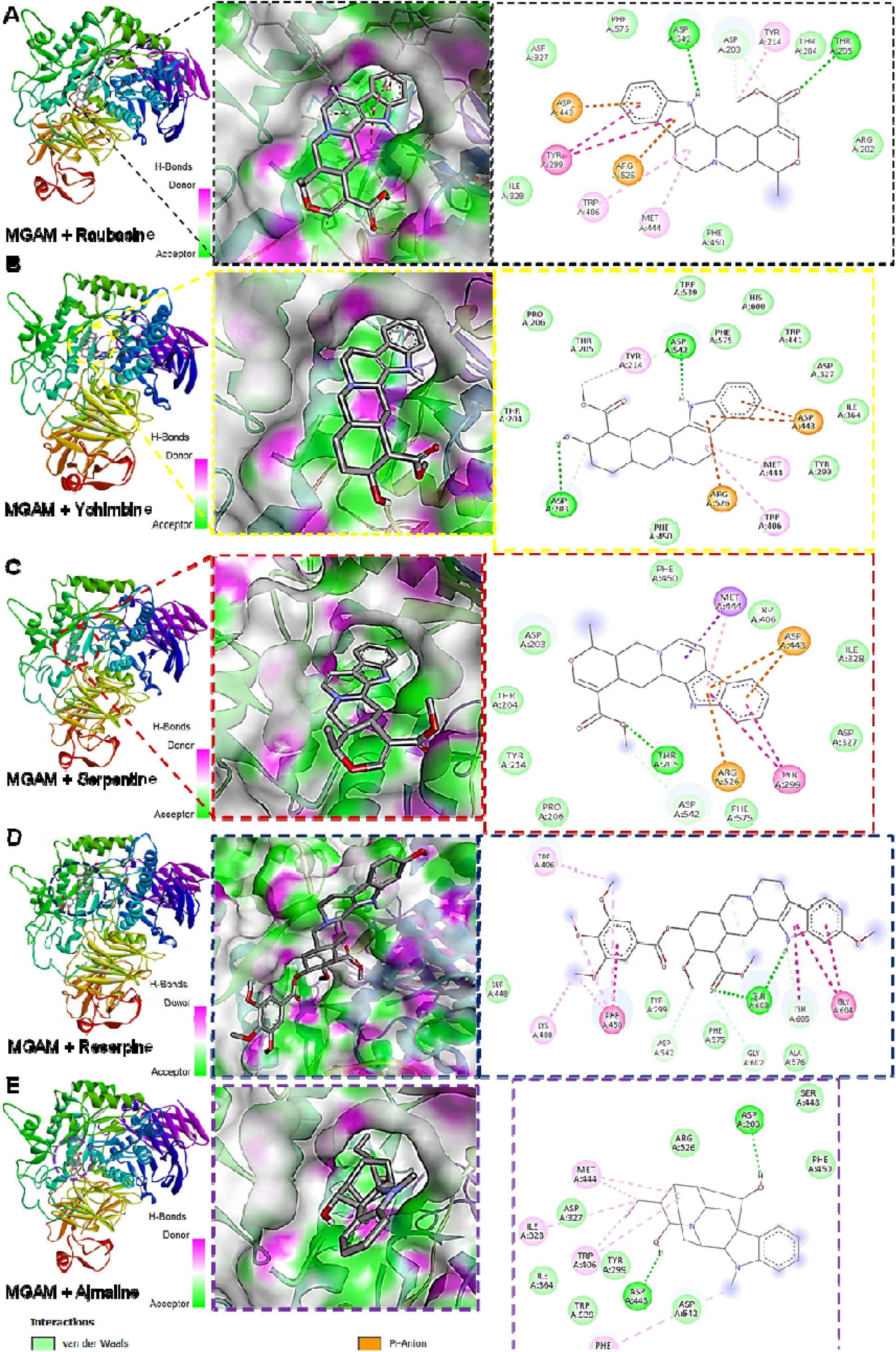

### In silico α-glucosidase inhibition activity of *Withania somnifera* phytochemicals

Molecular docking analysis was performed to evaluate the binding behavior of four major *Withania somnifera* phytochemicals including, Withaferin A, Withanone, Withanolide B, and Withanolide A within the catalytic pocket of MGAM. All compounds were favorably accommodated in the active site, exhibiting binding energies ranging from −6.0 to −9.25 kcal/mol, with distinct interaction patterns that reflect differences in binding strength and stability **(Figure 3A–D)**. Withaferin A showed moderate binding affinity toward MGAM with a binding energy of −6.0 kcal/mol **(Figure 3A)**. The ligand interacted with the active site through a combination of hydrogen bonding and hydrophobic contacts involving residues such as Asp443, Asp542, Tyr299, Trp406, Phe575, and Ile328, indicating stable but relatively weaker engagement of the catalytic pocket compared to other withanolides. Withanone demonstrated improved binding affinity (−7.57 kcal/mol) and formed hydrogen bond interactions with key catalytic residues including Asp542 and Thr205, supported by additional hydrophobic and π–π interactions with Tyr299, Trp406, Met444, and Phe575 **(Figure 3B)**. These interactions suggest enhanced stabilization within the MGAM active site relative to Withaferin A. Among the all *Withania somnifera* phytochemicals, Withanolide B exhibited the strongest binding affinity, with a binding energy of −9.25 kcal/mol **(Figure 3C)**. The ligand formed multiple hydrogen bonds with catalytically important residues such as Asp203, Asp443, Asp542, Gln603, and Arg526, while extensive hydrophobic and π–π interactions with Tyr299, Trp406, Met444, Phe575, and Ile328 further reinforced complex stability. This dense interaction network indicates tight binding and effective occupation of the MGAM catalytic pocket. Withanolide A also showed favorable binding with a binding energy of −7.35 kcal/mol **(Figure 3D)**. It interacted with MGAM through hydrogen bonds involving Asp203, Asp542, and Gln603, complemented by hydrophobic and aromatic interactions with Tyr299, Trp406, Phe450, Met444, and Ile364, suggesting stable active-site engagement. Overall, the docking results indicate that *Withania somnifera* phytochemicals, particularly Withanolide B and Withanone, exhibit strong binding affinities and well-distributed interaction networks within the MGAM catalytic pocket.

**Figure 3.**
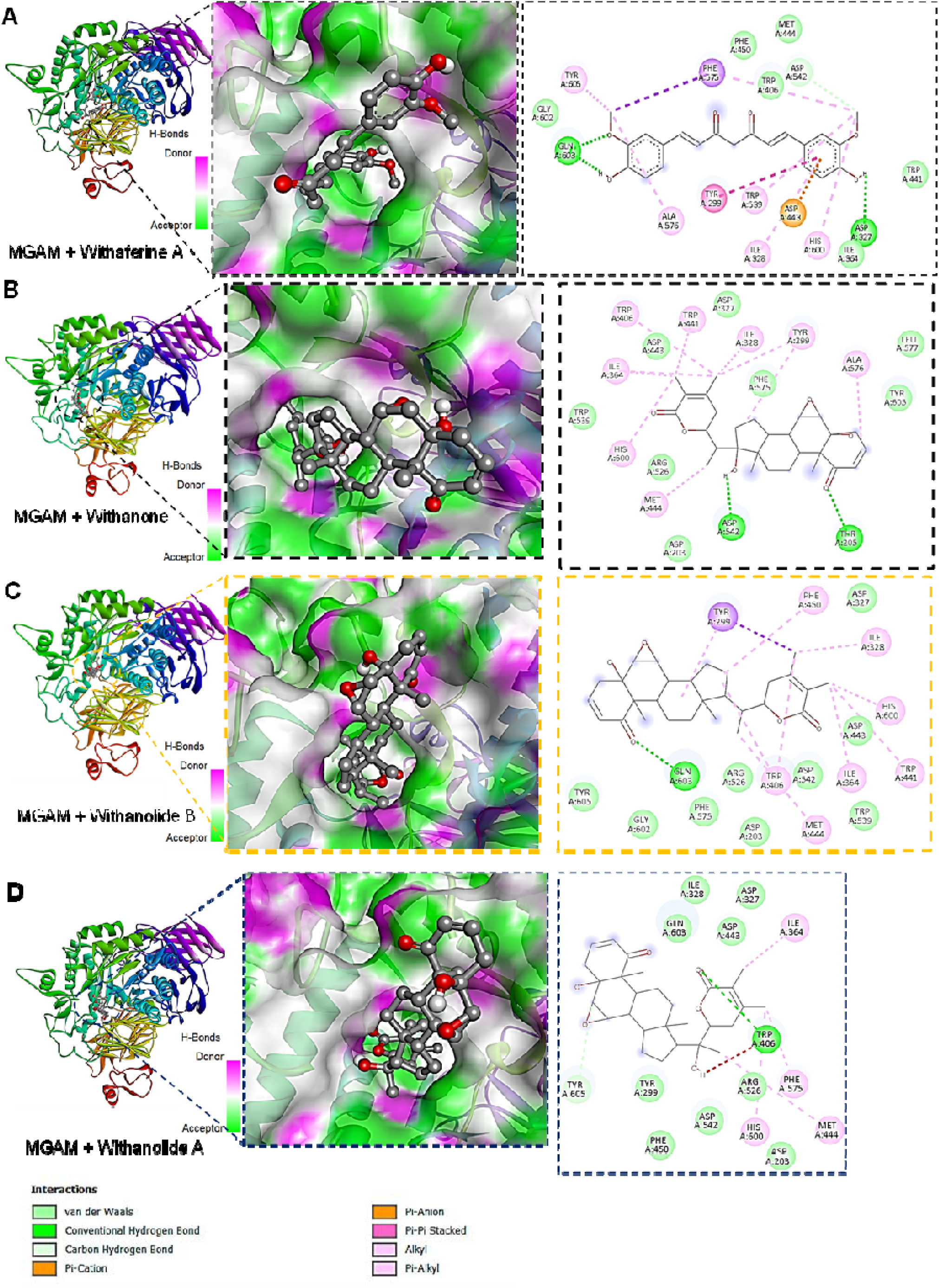

### In silico **α**-glucosidase inhibition activity of *Curcuma longa* phytochemicals

The binding behavior of major *Curcuma longa* phytochemicals, curcumin and deoxycurcumin, within the catalytic pocket of MGAM was evaluated using molecular docking analysis. Both compounds were favorably accommodated in the active site and exhibited comparable binding affinities, with binding energies of −6.36 kcal/mol for curcumin and −6.35 kcal/mol for deoxycurcumin, indicating moderate and stable interactions with MGAM **(Figure 4A and 4B)**. Curcumin showed stable binding within the MGAM active site with a binding energy of −6.36 kcal/mol **(Figure 4A)**. The ligand formed hydrogen bond interactions with key residues such as Asp327 and Gln603, which are involved in substrate recognition, while additional π–π and hydrophobic interactions with residues including Tyr299, Trp406, Phe575, Met444, and Ile328 contributed to overall complex stabilization. The extended conjugated structure of curcumin allowed effective alignment within the catalytic pocket, facilitating interactions across multiple regions of the active site. Deoxycurcumin exhibited a very similar binding profile, with a binding energy of −6.35 kcal/mol **(Figure 4B)**. It interacted with MGAM through hydrogen bonds involving Gln603 and Asp327, supported by hydrophobic and π–π interactions with Tyr299, Trp406, Met444, Phe450, and Ile364, indicating a comparable mode of active-site engagement to curcumin.

**Figure 4.**
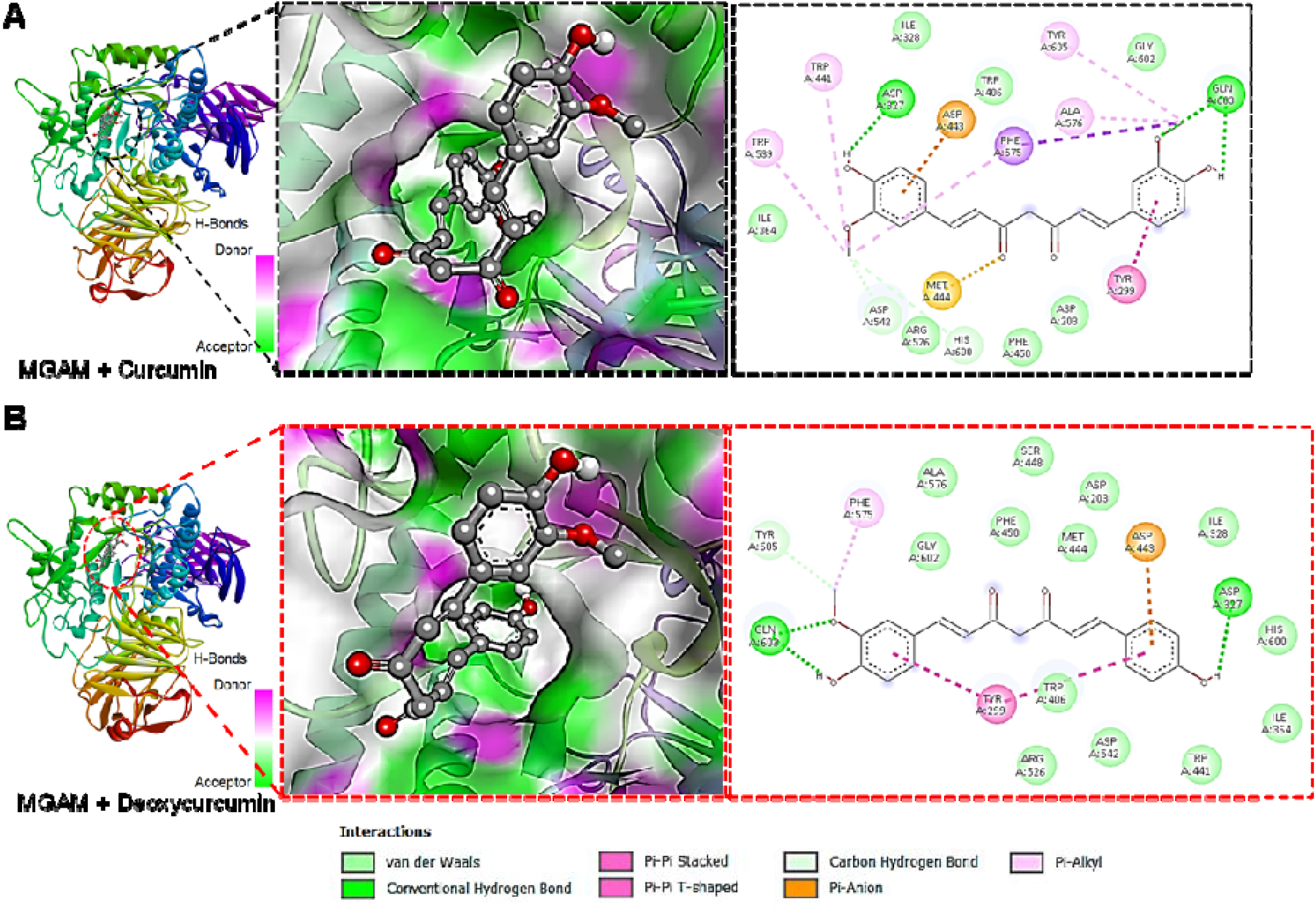

### In silico **α**-glucosidase inhibition activity of tea phytochemicals

Molecular docking analysis was performed to evaluate the interaction of major tea (*Camellia sinensis*) phytochemicals including, epicatechin gallate (ECG), epigallocatechin (EGC), epigallocatechin gallate (EGCG), and epicatechin with MGAM. All four compounds were successfully accommodated within the MGAM catalytic pocket and exhibited binding energies ranging from −5.65 to −6.85 kcal/mol, indicating moderate binding affinities with distinct interaction profiles **(Figure 5A–D)**. ECG displayed the strongest binding affinity among the tea phytochemicals, with a binding energy of −6.85 kcal/mol **(Figure 5A)**. ECG formed multiple hydrogen bonds with key catalytic residues, including Asp203, Asp327, Asp542, and Gln603, while additional π–π and hydrophobic interactions with residues such as Tyr299, Trp406, Phe575, Met444, and Ile328 contributed to stabilization of the ligand-enzyme complex. EGCG also showed favorable binding toward MGAM with a binding energy of −6.76 kcal/mol **(Figure 5C)**. The ligand established hydrogen bond interactions with catalytically important residues such as Asp203, Asp443, Asp542, and Gln603, supported by extensive hydrophobic and aromatic interactions involving Tyr299, Trp406, Met444, and Phe450, suggesting effective occupation of the active site. Epicatechin demonstrated moderate binding affinity with a binding energy of −6.15 kcal/mol **(Figure 5D)**. It interacted with MGAM through hydrogen bonds involving Asp203, Asp542, and Thr204, along with hydrophobic and π–π interactions with residues such as Tyr299, Trp406, Phe575, and Ile328, indicating stable but comparatively weaker active-site engagement. In contrast, EGC showed the lowest binding affinity among the tested tea phytochemicals, with a binding energy of −5.65 kcal/mol **(Figure 5B)**. Although EGC formed hydrogen bonds with residues including Asp203, Asp327, and Gln603 and maintained some hydrophobic interactions, the overall interaction network was less extensive, suggesting reduced stability within the MGAM catalytic pocket.

**Figure 5.**
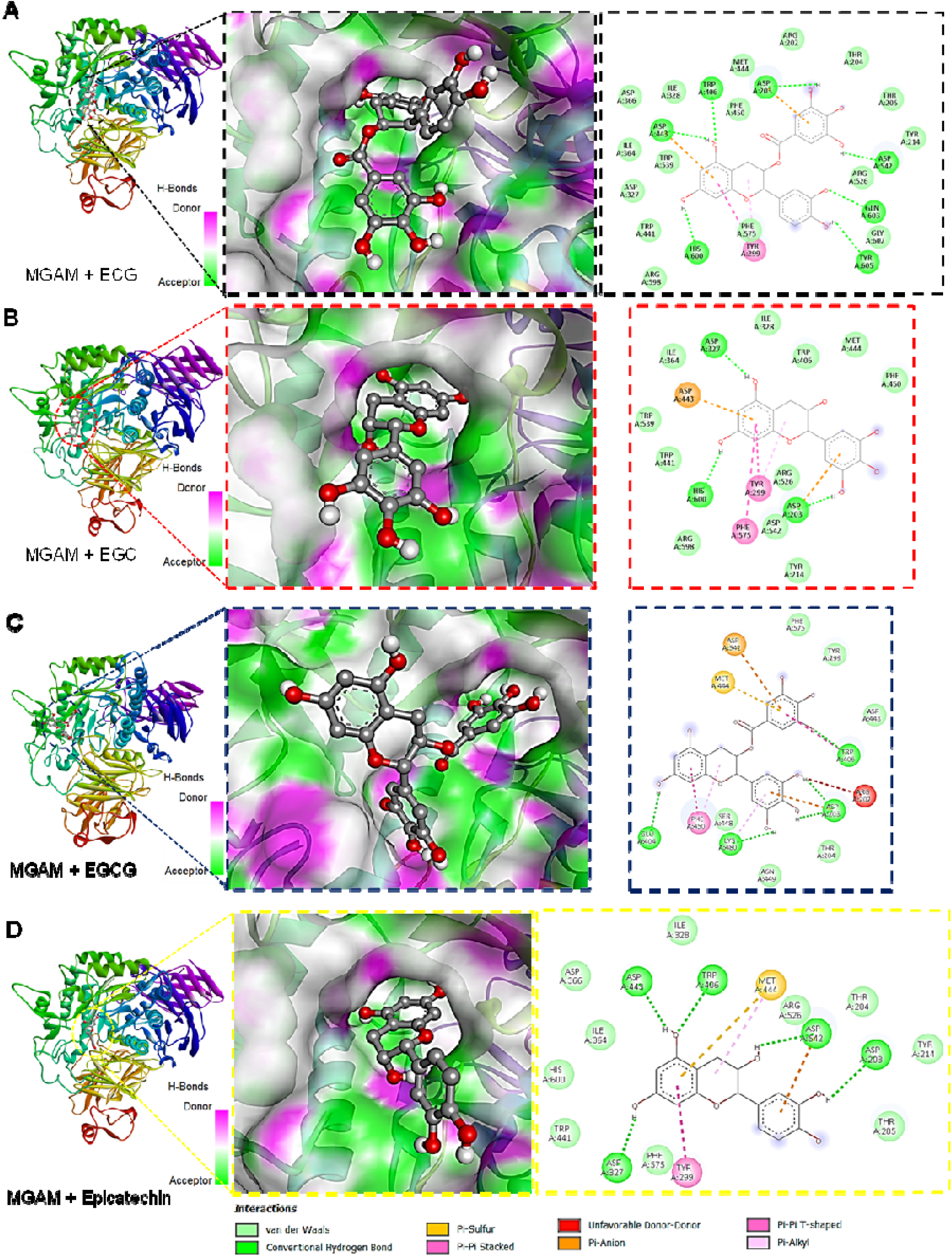

### ADME and drug-likeness evaluation of phytochemicals ADME profiling of the selected *R serpentine* phytochemicals

In silico ADME of yohimbine, raubasine, serpentine, ajmaline, and reserpine revealed distinct but generally favorable pharmacokinetic properties (**Figure 6 A-E and Supplementary file 1**). All compounds were predicted to have high gastrointestinal (GI) absorption, indicating significant potential for interaction at the intestinal interface. Solubility varied across the set, with ajmaline and serpentine showing good aqueous solubility, while yohimbine and raubasine were moderately soluble; resperine exhibited lower solubility predictions. Consensus lipophilicity (Log P) values were within acceptable ranges for most compounds, supporting balanced polarity and membrane interaction potential. Drug-likeness evaluations indicated full compliance with major filters (Lipinski, Ghose, Veber, Egan, Muegge) for yohimbine, raubasine, serpentine, and ajmaline, each also showing bioavailability scores of 0.55-0.85; in contrast, resperine displayed multiple violations and a low bioavailability score (0.17). P-glycoprotein substrate prediction was positive for yohimbine, raubasine, and ajmaline, while serpentine and resperine were not predicted to be P-gp substrates. None of the compounds were predicted to be significant inhibitors of major CYP450 enzymes (CYP1A2, CYP2C19, CYP2C9, CYP3A4), although some showed potential CYP2D6 inhibition. Detailed ADME profiles, including individual solubility classes, lipophilicity, drug-likeness parameters, and supplementary descriptors, are provided in **Supplementary File S1**.

**Figure 6.**
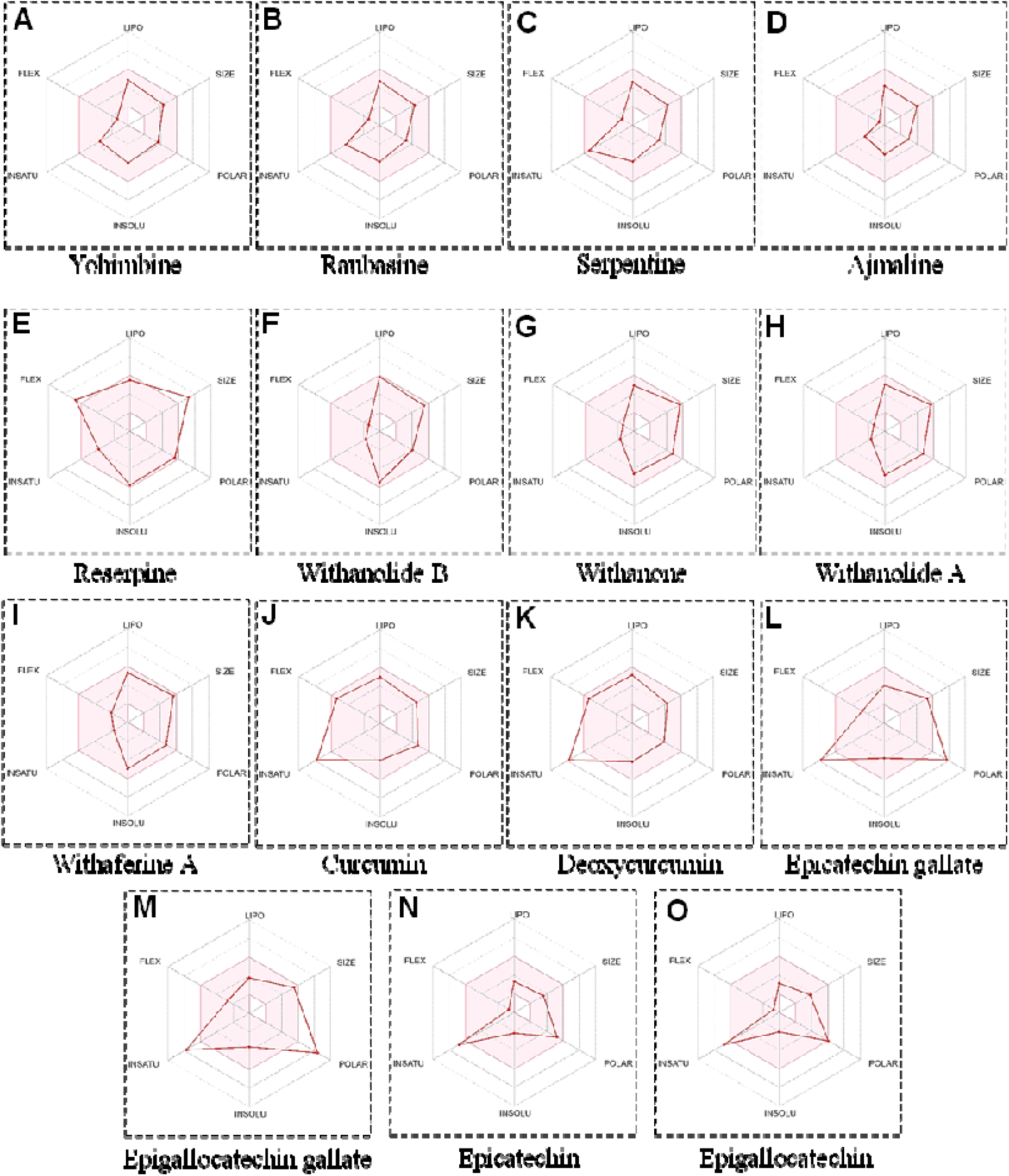

### ADME profiling of the selected *W. sominifera* phytochemicals

In silico ADME profiling of *Withania somnifera* phytochemicals revealed the following properties **(Figure 6 F-I and supplementary file 1)**. Withanolide B demonstrated high predicted gastrointestinal absorption and moderate aqueous solubility (Log S ESOL ∼ –5.46), balanced lipophilicity (consensus Log P ∼ 4.14), no blood–brain barrier permeation, and compliance with Lipinski and most drug-likeness rules; CYP2C9 inhibition was predicted. Withanone exhibited high predicted GI absorption, moderate solubility (Log S ESOL ∼ –4.55), consensus Log P ∼ 3.36, lack of BBB permeation, and favorable drug-likeness filters with one medicinal chemistry alert; no major CYP inhibition was predicted. Withanolide A showed high GI absorption, moderate solubility (Log S ESOL ∼ –4.67), consensus Log P ∼ 3.39, no BBB permeation, P-gp substrate prediction, compliance with major drug-likeness rules, and a single medicinal chemistry alert. Withaferine A was predicted with high GI absorption and moderate solubility (Log S ESOL ∼ –4.97), consensus Log P ∼ 3.42, no BBB permeation, P-gp substrate status, compliance with Lipinski and other filters, and one medicinal chemistry alert; all compounds had bioavailability scores of ∼ 0.55. Detailed ADME descriptors are provided in **Supplementary File S1**.

### ADME profiling of the selected *Curcuma longa* phytochemicals

In silico ADME profiling of *Curcuma longa* phytochemicals revealed the following characteristics (**Figure 6 J-K and supplementary file 1**). Curcumin exhibited high predicted gastrointestinal absorption with overall moderate aqueous solubility (Log S ESOL ≈ –3.94; SILICOS-IT ≈ –4.45), balanced lipophilicity (consensus Log P ≈ 3.03), and compliance with major drug-likeness filters (Lipinski, Ghose, Veber, Egan, Muegge), with a bioavailability score of 0.55; it was not predicted to permeate the blood–brain barrier, and showed no major CYP inhibition except CYP2C9. Deoxycurcumin also demonstrated high predicted GI absorption and moderate solubility (Log S ESOL ≈ –4.08; SILICOS-IT ≈ –5.04), acceptable lipophilicity (consensus Log P ≈ 3.35), and compliance with drug-likeness criteria with a bioavailability score of 0.55; it was not predicted to be a P-gp substrate and did not show significant inhibition of key CYP enzymes. Detailed ADME descriptors for each compound are provided in **Supplementary File S1**.

### ADME Prediction Results of Tea Phytochemicals

In silico ADME profiling results for the tea (*Camellia sinensis*) phytochemicals — epicatechin gallate (ECG), epigallocatechin gallate (EGCG), epicatechin, and epigallocatechin (EGC) — are summarized based on radar plots **(Figure 6 L-O)** with full profiles provided in **Supplementary File S1**. Epicatechin gallate (ECG) displayed low predicted gastrointestinal absorption, soluble to moderately soluble aqueous behavior (Log S ESOL ≈ –3.70 to –4.86), moderate consensus lipophilicity (∼1.30), and compliance with basic drug-likeness filters aside from a violation for hydrogen bond counts; no major CYP450 inhibition or P-gp substrate prediction was observed. Epigallocatechin gallate (EGCG) also showed low predicted GI absorption, good aqueous solubility (Log S ESOL ≈ –3.56 to –2.50), low to moderate lipophilicity (consensus Log P ≈ 0.95), and multiple drug-likeness rule violations primarily due to high polarity and hydrogen bond donors; no significant CYP450 inhibition or P-gp substrate status was predicted. Epicatechin exhibited high predicted GI absorption with good solubility (Log S ESOL ≈ –2.22 to –2.14), balanced lipophilicity (consensus Log P ≈ 0.85), and full compliance with Lipinski, Ghose, Veber, Egan, and Muegge filters; it was not predicted to inhibit major CYP enzymes or act as a P-gp substrate. Epigallocatechin (EGC) similarly showed high predicted GI absorption, good aqueous solubility (Log S ESOL ≈ –2.08 to –1.56), low lipophilicity (consensus Log P ≈ 0.42), and generally favorable drug-likeness except for a single Lipinski violation related to hydrogen bond donors; no significant CYP inhibition or P-gp substrate behavior was noted.

### Toxicity prediction of the phytochemicals

The oral toxicity of the selected phytochemicals and the standard inhibitor miglitol was predicted using in silico approaches to estimate acute toxicity risk following ingestion. Predicted LD values (mg/kg) and corresponding toxicity classes are presented in Table 2. Across the evaluated compounds, Class 2 and Class 3 toxicities were observed for several alkaloids and withanolides, indicating moderate acute toxicity at high systemic doses. Among *Rauwolfia serpentina* phytochemicals, yohimbine, raubasine, serpentine, and reserpine were predicted with LD values of approximately 215–300 mg/kg (Class 3), indicating moderate predicted acute toxicity, whereas ajmaline showed a lower LD (∼34 mg/kg, Class 2), suggesting higher acute toxicity if absorbed systemically. Similarly, *Withania somnifera* withanolides such as withanolide B, withanone, and withanolide A were predicted to have LD values in the range of 7–34 mg/kg (Class 2), pointing to a higher systemic toxicity potential when compared with Class 3 compounds, while withaferin A exhibited a predicted LD of ∼300 mg/kg (Class 3). In contrast, phytochemicals from *Curcuma longa* (curcumin and deoxycurcumin) and *Camellia sinensis* (ECG, EGCG, epicatechin, and EGC) generally demonstrated higher LD values (1000–10000 mg/kg; Class 4–6), indicating lower predicted acute toxicity. The standard inhibitor miglitol was predicted with an LD of ∼1200 mg/kg (Class 4), consistent with low acute oral toxicity.

**Table 1.** Binding energies of phytochemicals from *Rauwolfia serpentina*, Withania somnifera, Curcuma longa, and *Camellia sinensis* docked against maltase-glucoamylase (MGAM), ranked by binding affinity.

| Plant source | Compound name | Binding energy (kcal/mol) | Rank (within plant) |
| --- | --- | --- | --- |
| *Rauwolfia serpentina* | Yohimbine | **-8.50** | 1 |
|  | Raubasine | -8.46 | 2 |
|  | Serpentine | -7.62 | 3 |
|  | Ajmaline | -7.53 | 4 |
|  | Reserpine | -6.68 | 5 |
| *Withania somnifera* | Withanolide B | **-9.25** | 1 |
|  | Withanone | -7.57 | 2 |
|  | Withanolide A | -7.35 | 3 |
|  | Withaferin A | -6.00 | 4 |
| *Curcuma longa* | Curcumin | **-6.36** | 1 |
|  | Deoxycurcumin | -6.35 | 2 |
| *Camellia sinensis* | Epicatechin gallate (ECG) | **-6.85** | 1 |
|  | Epigallocatechin gallate (EGCG) | -6.76 | 2 |
|  | Epicatechin | -6.15 | 3 |
|  | Epigallocatechin (EGC) | -5.65 | 4 |
| Standard inhibitor | Miglitol | -6.86 | - |

**Table 2:**
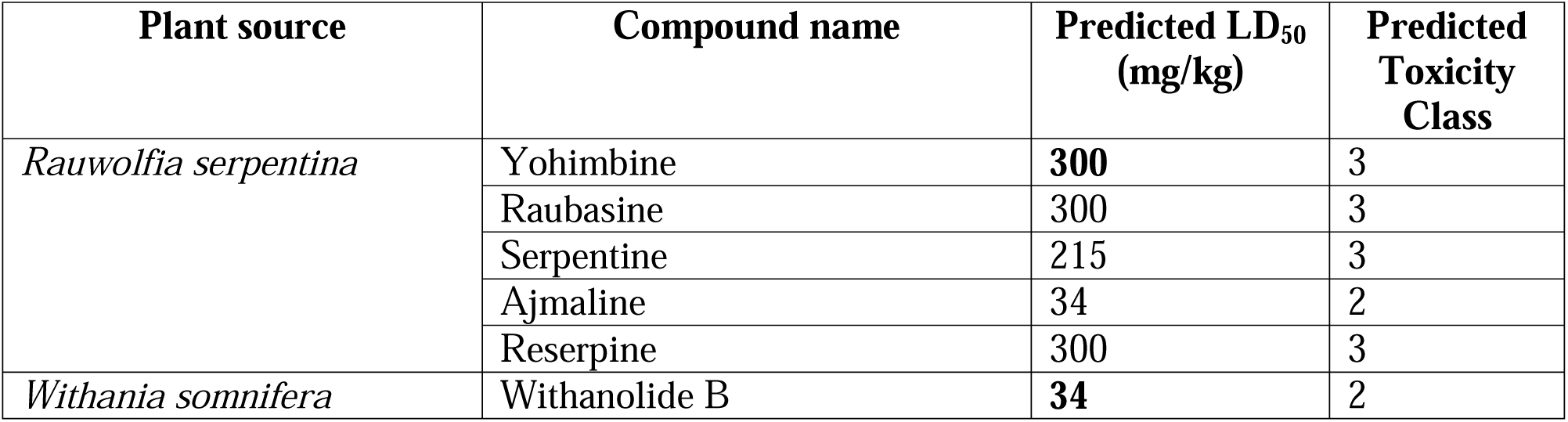
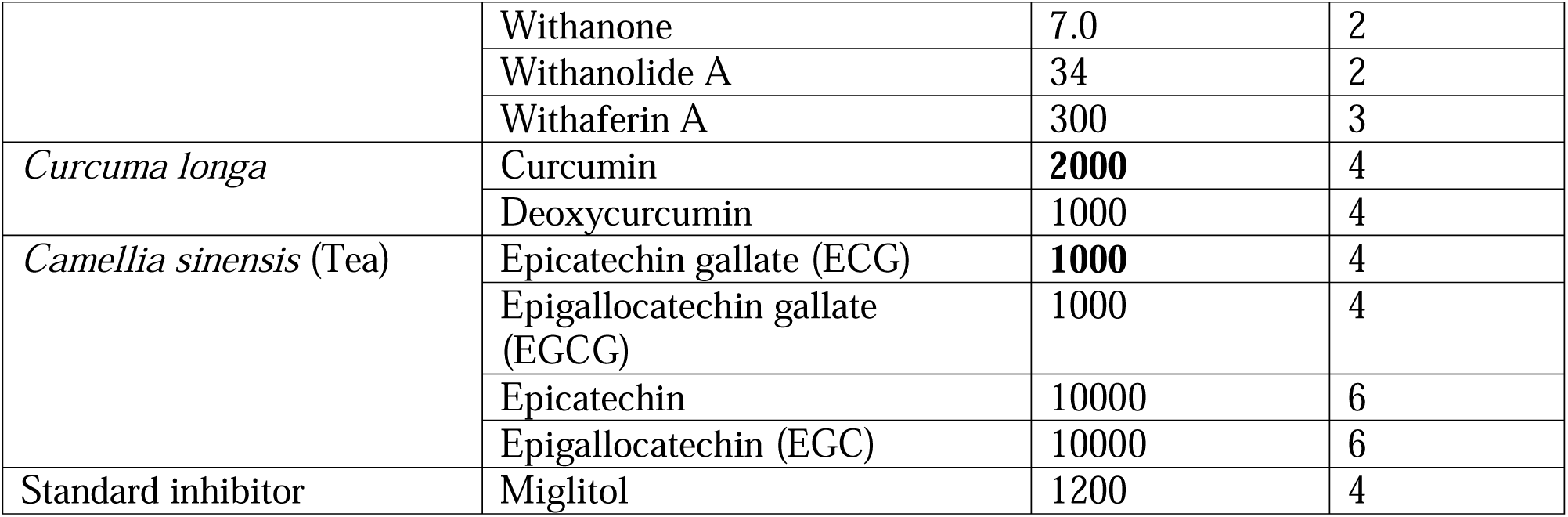
Toxicity profiling of the phytochemicals using ProTox database.

## Discussion

Postprandial hyperglycemia plays a central role in the pathophysiology of type 2 diabetes mellitus (T2DM), contributing to cardiovascular and microvascular complications. Delaying or reducing the rate of carbohydrate digestion in the intestine is an established therapeutic strategy to control postprandial glucose excursions (Zhang et al. 2021). The small-intestinal brush border houses a family of α-glucosidases, chiefly maltase–glucoamylase (MGAM) and sucrase-isomaltase, which catalyze the final hydrolysis of starch-derived oligosaccharides to glucose, facilitating absorption across enterocytes (Rose et al. 2018). Inhibiting these enzymes can slow the conversion of dietary starch into absorbable glucose, reducing the magnitude of postprandial plasma glucose peaks – a mechanism exploited by clinical α-glucosidase inhibitors such as miglitol, acarbose, and voglibose (Alssema et al. 2021) Miglitol, a synthetic α-glucosidase inhibitor widely used in T2DM, binds to MGAM’s catalytic region predominantly through multiple polar hydrogen bonds, mimicking glucose and thereby competitively inhibiting enzyme activity (Akmal et al. 2025). In this study, we evaluated the potential of phytochemicals from Withania somnifera, *Rauwolfia serpentina*, Curcuma longa, and *Camellia sinensis* to inhibit maltase–glucoamylase (MGAM), using the standard α-glucosidase inhibitor miglitol as a comparative benchmark to assess relative binding affinities and interaction profiles. The docking results indicate that *Withania somnifera* phytochemicals, especially Withanolide B and Withanone, exhibited the most robust predicted interactions with the MGAM catalytic site.

These compounds not only formed multiple hydrogen bonds with key residues of MGAM but also established stabilizing hydrophobic and π–π interactions, which may enhance conformational stability within the active site. Withanolides are a class of steroidal lactones recognized as major bioactive constituents of W. somnifera, traditionally used as a Rasayana herb in Ayurveda for managing metabolic disorders, including diabetes (Akanksha et al. 2022; Mirjalili et al. 2009). Systematic reviews suggest that *Withania somnifera* treatment significantly restores altered blood glucose and glycosylated hemoglobin (HbA1c) levels and improves metabolic profiles in diabetic models with minimal safety concerns (Durg et al. 2020; Udayakumar et al. 2009). Animal studies corroborate the antidiabetic potential of W. somnifera, showing hypoglycemic and hypolipidemic effects in alloxan-induced diabetic rats (Udayakumar et al. 2009) and improvements in insulin sensitivity accompanied by reduced inflammatory markers in fructose-fed rats (Samadi Noshahr et al. 2015). In silico ADME profiling predicted that major *Withania somnifera* phytochemicals such as Withanolide B, Withanone, Withanolide A, and Withaferin A have high gastrointestinal absorption, moderate aqueous solubility, balanced lipophilicity, and generally comply with major drug-likeness rules (e.g., Lipinski), while none are predicted to permeate the blood–brain barrier. These properties suggest favorable intestinal bioavailability and oral exposure for these compounds, supporting their potential as orally acting MGAM inhibitors. Empirical human pharmacokinetic data confirm that withanolides from *Withania somnifera* extracts are absorbed following oral administration, as evidenced by measurable plasma concentrations and pharmacokinetic parameters by (Rathi and Kim 2025).

However, in silico toxicity predictions indicated Class 2–3 toxicity levels for withanolides; actual toxicity may differ in biological scenarios. Moreover, the extensive traditional use of *Withania somnifera* and multiple experimental toxicity evaluations of its root and whole-plant extracts have generally demonstrated a favorable safety profile, with high tolerated doses in repeated-dose animal studies and an absence of significant adverse effects in human clinical evaluations. A 28-day subacute toxicity evaluation under OECD-guideline conditions demonstrated that *Withania somnifera* whole-plant extract exhibited no significant adverse effects in rats, supporting the acceptability of long-term use (Langade et al. 2023).Clinical studies in healthy adult volunteers indicate that *Withania somnifera* (*Withania somnifera*) root extract is generally well tolerated when taken orally at commonly studied dosages. An 8-week randomized, placebo-controlled trial showed that supplementation with *Withania somnifera* root extract was safe in both male and female participants, with no significant adverse events reported throughout the study period (Verma et al. 2021). Additional clinical evaluations have demonstrated that healthy subjects can tolerate standardized *Withania somnifera* supplementation (e.g., up to 1000 mg daily) for several weeks without observable safety concerns in physiological or biochemical parameters (Vaidya et al. 2024).

Although miglitol, the standard inhibitor for MGAM, effectively lowers postprandial glucose levels, its interaction profile lacks extensive hydrophobic stabilization within the MGAM pocket when compared with several *Withania somnifera* phytochemicals examined in this study, which form additional π–π and hydrophobic contacts. This distinction in interaction type may partly explain the stronger binding energies seen in silico for certain plant-derived ligands with MGAM. Therefore, *Withania somnifera* extract may be a potential option for helping maintain postprandial glucose levels in diabetic patients, although further experimental validation is required.

After *Withania somnifera* phytochemicals, *Rauwolfia serpentina* phytochemicals exhibited strong binding affinity toward MGAM, with binding energies ranging from −8.50 to −6.68 kcal/mol; among them, yohimbine was identified as the most effective compound. However, yohimbine has been reported to induce systemic toxicity in different organs (Bourgeois and Eggleston 2023; Giampreti et al. 2009; Kearney et al. 2010), although it is predicted to be less toxic than withanolides. R. serpentina has a long history in traditional medicine for cardiovascular and metabolic conditions (Azmi and Qureshi 2012). Despite its therapeutic promise, *Rauwolfia serpentina* and its principal alkaloids (e.g., reserpine) are associated with systemic adverse effects such as gastrointestinal disturbances, hypotension, depressive symptoms, and sexual dysfunction — risks that underscore the need for careful pharmacological and toxicological evaluation before clinical application (Khan et al. 2025).

Curcuminoids from *Curcuma longa* (curcumin and deoxycurcumin) and tea phytochemicals showed moderate MGAM binding energies, comparable to those of some other plant bioactives, but exhibited a safer toxicological profile. But in combination of MGAM binding cability and toxicological profile, *Withania somnifera* extract enriched with withanolide A& B, withanone will be agood target for managing postparandiral blood glucose. By preventing or slowing postprandial surges in blood sugar, these agents can improve overall glycemic control, potentially reducing insulin demand and mitigating long-term complications associated with chronic hyperglycemia.

Further, standardized *Withania somnifera* extracts enriched for withanolides, particularly Withanolide B and Withanone, should be evaluated using purified enzyme assays and animal glucose tolerance models. Adjunctive agents like curcuminoids and tea catechins may also provide incremental benefits as part of multifactorial dietary interventions, although their direct impact on MGAM likely requires enhancement through targeted delivery.

### Conclusion

In conclusion, in silico evaluation revealed that *Withania somnifera* phytochemicals, particularly Withanolide B and Withanone, showed the strongest predicted binding to the MGAM active site, suggesting enhanced potential to slow carbohydrate digestion and attenuate postprandial glucose spikes. These compounds formed multiple stabilizing interactions within the MGAM catalytic pocket and have predicted properties supportive of oral intestinal exposure. Compared with miglitol, the plant-derived ligands exhibited additional hydrophobic and π–π contacts, which may contribute to comparatively stronger binding energies in silico. While other botanicals demonstrated moderate MGAM affinity or safety concerns, the withanolide-rich *Withania somnifera* extract emerged as the most promising candidate for helping moderate postprandial glucose excursions. Therefore, standardized extracts enriched in key withanolides warrant further experimental validation in enzyme assays and glucose tolerance models.

## CRediT authorship contribution statement

**Ashim Chandra Roy:** Conceptualization, Methodology, Investigation, Resources, Data curation, Writing – original draft, Review & Editing, Validation, Formal analysis, Visualisation, Software. **Ilora Ghosh:** Conceptualization, Resources, Writing – review & editing, Validation, Supervision.

## Data Availability

Data will be made available on request.

## Acknowledgements

ACR thanks DBT (Government of India) and KP thanks CSIR (Government of India), for providing Senior Research Fellowships. The funding agencies had no role in study design, data acquisition, analysis, the decision to publish, and preparation of the manuscript.

## Consent to participate

Not applicable

## Consent to publish

Not applicable

